# Integration of eQTL and Parkinson’s disease GWAS data implicates 11 disease genes

**DOI:** 10.1101/627216

**Authors:** Demis A. Kia, David Zhang, Sebastian Guelfi, Claudia Manzoni, Leon Hubbard, United Kingdom Brain Expression Consortium (UKBEC), International Parkinson’s Disease Genomics Consortium (IPDGC), Regina H. Reynolds, Juan Botía, Mina Ryten, Raffaele Ferrari, Patrick A. Lewis, Nigel Williams, Daniah Trabzuni, John Hardy, Nicholas W. Wood

**Affiliations:** Department of Molecular Neuroscience, UCL Institute of Neurology, Queen Square, London WC1N 3BG, UK; School of Pharmacy, University of Reading, Reading RG6 6AP, UK; The complete list of the IPDGC and UKBEC members is listed in the Supplementary Material; Departamento de Ingeniería de la Información y las Comunicaciones. Universidad de Murcia, Murcia, Spain; Department of Genetics, King Faisal Specialist Hospital and Research Centre, 11211 Riyadh, Saudi Arabia

**Keywords:** Human brain expression and splicing QTLs, Parkinson’s disease (PD) GWAS, COLOC, TWAS, *ZRANB3*, *PCGF3*, *NEK1*, *NUPL2*, *GALC*, *CTSB*, *WDR6*, *CD38*, *GPNMB*, *RAB29*, *TMEM163*

## Abstract

Substantial genome-wide association study (GWAS) work in Parkinson’s disease (PD) has led to an increasing number of loci shown reliably and robustly to be associated with the increased risk of the disease. Prioritising causative genes and pathways from these studies has proven problematic. Here, we present a comprehensive analysis of PD GWAS data with expression and methylation quantitative trait loci (eQTL/mQTL) using Colocalisation analysis (Coloc) and transcriptome-wide association analysis (TWAS) to uncover putative gene expression and splicing mechanisms driving PD GWAS signals. Candidate genes were further characterised by determining cell-type specificity, weighted gene co-expression (WGNCA) and protein-protein interaction (PPI) networks.

Gene-level analysis of expression revealed 5 genes (*WDR6, CD38, GPNMB, RAB29, TMEM163*) that replicated using both Coloc and TWAS analyses in both GTEx and Braineac expression datasets. A further 6 genes (*ZRANB3, PCGF3, NEK1, NUPL2, GALC, CTSB*) showed evidence of disease-associated splicing effects. Cell-type specificity analysis revealed that gene expression was overall more prevalent in glial cell-types compared to neurons. The WGNCA analysis showed that *NUPL2* is a key gene in 3 modules implicated in catabolic processes related with protein ubiquitination (protein ubiquitination (p=7.47e-10) and ubiquitin-dependent protein catabolic process (p = 2.57e-17) in *nucleus accumbens, caudate* and *putamen*, while *TMEM163* and *ZRANB3* were both important in modules indicating regulation of signalling (p=1.33e-65] and cell communication (p=7.55e-35) in the *frontal cortex* and *caudate* respectively. PPI analysis and simulations using random networks demonstrated that the candidate genes interact significantly more with known Mendelian PD and parkinsonism proteins than would be expected by chance. The proteins core proteins this network were enriched for regulation of the ERBB receptor tyrosine protein kinase signalling pathways.

Together, these results point to a number of candidate genes and pathways that are driving the associations observed in PD GWAS studies.

## Introduction

Parkinson’s disease (PD) is the second most common neurodegenerative condition worldwide, characterised by bradykinesia, rigidity and tremor (Lees *et al.*). Recent efforts in genome-wide associated studies have unveiled over 40 risk loci (Chang *et al.*, 2017a). However, the genes and the genomic processes driving the disease mechanisms remain largely unknown. Most genome wide associations studies (GWAS) risk variants fall in the non-coding regions of the genome. Several studies show that complex trait-associated variants can act as expression and splicing quantitative trait loci (eQTL/sQTL), and exploring eQTLs has proven a fruitful method to follow up GWAS results (Emilsson *et al.*, 2008; Nica *et al.*, 2010). An improved understanding of the underlying mechanisms via which these risk variants act will be instrumental to the understanding of PD pathophysiology.

Several public brain tissue-specific eQTL datasets have become available in recent years. These include the Braineac dataset by the UK Brain Expression Consortium (UKBEC) (Trabzuni *et al.*, 2011a) and the Genotype-Tissue Expression (GTEx) consortium (Ramasamy *et al.*, 2014; The GTEx Consortium, 2015). Such datasets have enabled the interpretation of GWAS results in the context of regulatory effects of risk variants on genes. These datasets offer differing opportunities and limitations, CommonMind has a large number but only one brain region, whereas Gtex and Braineac provide overlapping but non-identical brain regions and provide a platform for splicing and regulatory analyses.

There have also been recent advances in statistical methods allowing much more detailed interrogation of GWAS and eQTL summary association data to elucidate complex disease mechanisms: Coloc (Giambartolomei *et al.*, 2014) is a method that uses a Bayesian framework to assign a posterior probability to the hypothesis that two traits share a causal variant. Other methods quantify the genetic correlation between gene expression and disease GWAS; Summary Mendelian randomization (SMR) leverages Mendelian randomization methods to assess the degree to which the disease outcome could be causally explained by changes in gene expression (Zhu *et al.*, 2016); Transcriptome-wide association study (TWAS) and MetaXcan are based on marrying disease GWAS data with prediction models trained on reference expression data to assess the association between gene expression and disease (Gusev *et al.*, 2016; Barbeira *et al.*, 2017). Here we have selected Coloc and TWAS as the primary methods of eQTL analysis, in order to gain complementary information on the relationship between gene expression and PD (Gusev, 2017).

The availability of large transcriptomic datasets and advanced statistical tools and methods to integrate expression platform with GWAS data improves our ability to identify candidate disease-causing genes to investigate further (Dobbyn *et al.*, 2017; Pardiñas *et al.*, 2018). Recent efforts in PD at the 7p15.3 locus have shown the risk variants in *GPNMB* act as expression QTL (Murthy *et al.*, 2017).

Here we present a systematic interrogation of PD risk loci from the most recent GWAS (Chang *et al.*, 2017b) to uncover putative genes and genomic events, within the PD risk loci, based on gene expression, splicing regulation and methylation in the human brain. This work is a reference example to be applied to different neurodegenerative diseases.

## Methods

### Parkinson’s disease GWAS data

Summary statistics from the combined discovery and replication phases of the recent GWAS meta-analysis of PD were used (Chang *et al.*, 2017b). This includes 8,055,803 genotyped and imputed variants in up to 26,035 PD cases and 403,190 controls of European ancestry. For the purposes of this study, all alleles were aligned on the forward strand, and all effect sizes and allele frequencies were converted with respect to the non-reference allele in build GRCh37. All genes overlapping the region 1Mb up or downstream of a SNP with a PD p-value ≤ 5×10^-8^ were selected for the initial analysis. The analysis was then extended to include all genes in the genome, to identify candidate genes in loci that have not reached genome-wide significance in the PD GWAS, but where the collective evidence with expression data suggests a colocalised signal.

### Braineac eQTL data

The UKBEC Braineac dataset contains data from 10 brain regions obtained from 134 control individuals: frontal cortex, temporal cortex, occipital cortex, hippocampus, thalamus, putamen, substantia nigra, medulla, cerebellum, and white matter, together with the average expression across all 10 regions (Trabzuni *et al.*, 2011b; Trabzuni *et al.*, 2013; Ramasamy *et al.*, 2014). Gene expression was quantified using Affymetrix Exon 1.0 ST arrays, and the genotyping was performed using Illumina Infinium Human Omni1-Quad BeadChip microarrays then imputed to the European panel of the phase-1 1000 Genomes Project (Li *et al.*, 2009; Li *et al.*, 2010). The genotyped and imputed data were restricted to ∼5.88 million SNPs with MAF ≥ 0.05 and imputation r^2^ > 0.5. For each gene of interest, all SNP associations within 1Mb up and downstream of the gene were collected (http://www.braineac.org).

### GTEx eQTL data

The GTEx V7 dataset (The GTEx Consortium, 2015) includes eQTL data from 13 brain tissues with sample sizes ranging from 154 to 80: cerebellum, caudate, cortex, nucleus accumbens, cerebellar hemisphere, frontal cortex, putamen, hippocampus, anterior cingulate cortex, hypothalamus, amygdala, spinal cord (cervical C1), substantia nigra. Gene expression in these samples has been obtained using paired-end RNA-seq (Illumina TruSeq), and genotype data from whole genome sequencing. Full summary eQTL data for the tissues of interest was downloaded from the GTEx web portal (http://www.gtexportal.org/home/).

### Brain methylation data

Genome-wide methylation profiles were obtained from both substantia nigra and frontal cortex of 134 individuals with PD from the Parkinson’s Disease UK Brain Bank, using the Illumina Infinium HumanMethylation450 BeadChip (HM450). Cis PDmQTLs were defined as correlations between the target PD SNP genotype and DNAm levels of CpGs within a 500kb window of the SNP base position. Linear models were fitted to test whether DNAm beta-values for each CpG were predicted by SNP genotypes. We included covariates for age at death, gender, population stratification, batch and post-mortem interval. We retained the strongest SNP-CpG pair at a 5% false discovery rate to be used in downstream analyses. CpGs were mapped to genes if they were within 10kb of the gene transcription start / end base position according to HG19 coordinates.

### Coloc analysis

To assess the probability of the same SNP being responsible for both change in PD risk and modulating the expression levels of a gene, we used the Coloc method (Giambartolomei *et al.*, 2014). Both Braineac and GTEx eQTL datasets were harmonised with the PD GWAS dataset, to ensure that the regression coefficients were reported with respect to the non-reference alleles in build GRCh37, and variants overlapping with the PD-GWAS dataset were kept for analysis. Coloc uses estimated approximate Bayes factors from summary association data to compute posterior probabilities for five hypotheses:

**H_0_** No shared causal variant in the region

**H_1_** There is a causal PD variant but no eQTL variant

**H_2_** There is a causal eQTL variant but no PD variant

**H_3_** Both studies have a different causal variant within the analysed region

**H_4_** There is a shared causal variant within the analysed region

We used the default Coloc priors of *p*_*1*_=10^-4^, *p*_*2*_=10^-4^, *p*_*12*_=10^-5^ (*p*_*1*_ = the probability that a given SNP is associated with PD, *p*_*2*_ = probability that a given SNP is a significant eQTL, *p*_*12*_ = probability that a given SNP is both a PD hit and an eQTL).

For both the Braineac and GTEx datasets, we derived posterior probabilities (PPH0-4) for each gene and considered PPH4 ≥ 0.75 as strong evidence for colocalization. For Braineac, we also looked at genes where there is strong evidence of colocalization at exon-level for a given exon (exon PPH4 ≥ 0.75), but evidence against colocalization for the whole gene (gene PPH3 > gene PPH4), to identify potential splicing events causing PD.

### TWAS

To assess the degree to which changes in gene expression or splicing might be associated with PD case/control status, we performed a TWAS/MWAS using the method by Gusev et al. (Gusev *et al.*, 2016). Expression reference weights were obtained from the CommonMind Consortium (CMC) dorsolateral prefrontal cortex (DLPFC) RNA-seq and RNA-seq splicing datasets, which are based on 467 samples (209 schizophrenia cases, 206 controls, 52 affective disorder cases), and methylation data from our PD brain methylation dataset (Senthil *et al.*, 2017). For all genes (or isoforms for splicing analysis), TWAS/MWAS p-values were obtained, and all genes/isoforms passing multiple testing correction at FDR 0.05 level, genome-wide were considered as significant. Where multiple genes were implicated within a region, we performed further conditional analyses using Fusion to identify whether there were single or joint TWAS/MWAS signals at each locus. Conditional analyses were performed independently across gene expression and methylation datasets.

### Weighted gene co-expression network analysis (WGCNA)

We performed a weighted gene co-expression network analysis (WGCNA) with k-means applied to transcriptomic data from GTEx and Braineac to generate co-expression modules (Forabosco *et al.*, 2013; Bettencourt *et al.*, 2014; Botía *et al.*, 2017). Briefly, each module is associated to a cell type based on the enrichment of cell type-specific genes within the module. The enrichment is assessed by using a Fisher’s exact test to determine whether we find an overlap between the module genes and the brain cell type markers that is more significant than random chance. Each gene of interest is then assigned to a “primary” cell type based on its module membership (MM). Module membership is the correlation of the expression of our gene of interest with the first principal component of each module. This correlation is always between 0 and 1And we use MM as a measure of how reliable is the assignment of each gene to its module.

### Cell-type specificity analysis

We investigated the cell-type specific expression of the Coloc prioritised genes, using the immunopanning data from humans and mouse, and co-expression analysis of the GTEx and Braineac data (Zhang *et al.*, 2014; Zhang *et al.*, 2016; Soreq *et al.*, 2017). From the immunopanning data, cell type-specific enrichment values were obtained or calculated for each gene and for each cell type analysed. Enrichment was calculated as expression prevalence by dividing the average expression of the gene in one cell type by the average expression across all other cell types. Each gene of interest was then assigned to a “primary cell type” of interest based on the highest cell type-specific enrichment value observed.

### Literature-derived PPI networks

We extracted currently known protein interactors (PPIs) for the proteins (seeds) encoded by the genes prioritized in this manuscript (coloc protein network). PPIs were identified for each seed protein based upon entries in the following databases within the IMEX consortium (PMID: 22453911): APID Interactomes, BioGrid, bhf-ucl, InnateDB, InnateDB-All, IntAct, mentha, MINT, InnateDB-IMEx, UniProt, and MBInfo by means of the “PSICQUIC” R package (version 1.15.0 by Paul Shannon, http://code.google.com/p/psicquic/). For 2 of the seeds (TMEM163 and NEK) no human protein interaction data was available thus they were excluded form the analysis. After downloading PPIs (21 January 2018), Protein IDs were converted to uniprot and Entrez IDs. All databases were merged after removal of TrEMBL, non-protein interactors (e.g. chemicals), obsolete Entrez and Entrez matching to multiple Swiss-Prot identifiers. All PPIs underwent quality control (QC) following the weighted protein-protein interaction network analysis (WPPINA) pipeline as previously described to remove: i) all non-human annotations, and; ii) all poor quality annotations (Ferrari *et al.*, 2017). The interactions were then scored taking into consideration the number of different publications reporting the interaction, and the number of different methods reporting the interaction. All the interactors with a final score ≤ 2 were discarded to control for replication and reduce false positive rate. Of note, UBC (coding for the protein ubiquitin) was removed due to the pervasive nature of covalent ubiquitylation of proteins as part of the ubiquitin/proteasome degradation pathway. A similar network (Mendelian protein network) was prepared for Mendelian, Parkinson’s/parkinsonism proteins (*SNCA, LRRK2, GBA, SMPD1, VPS35, DNAJC13, PINK1, PRKN, DJ1, FBXO7, SYNJ1, DNAJC6, PRKRA, C19Orf12, PANK2, SPG11, RAB39B, ATP13A2, PLA2G6* and *WDR45*) and for 118 genes with a negative score in the coloc analysis (the latter being used as a negative control protein network, **Supplementary Table 1**. The final networks were visualized through the freely available Cytoscape 3.5.0 software (Shannon *et al.*, 2003). Functional enrichment was on the relevant genes prioritized through WPPINA was performed using g:Profiler (doi: 10.1093/nar/gkw199) on the 8th February 2018 allowing for Gene Ontology (GO) Biological Processes (BP) terms to be queried.

## Results

### Gene-level results

Overall, 515 genes were present within 1Mb of a genome-wide significant PD SNP in the Braineac dataset, and 748 in at least one GTEx brain tissue dataset, with 470 of these overlapping. In Braineac, 9 genes had strong evidence for colocalization at gene level in at least one brain region, while in GTEx, 42 genes had strong evidence for association. Fifteen of these 42 genes were not present in Braineac, so replication across both datasets was possible in 27 genes. In the TWAS analysis, 137 genes were significant at FDR 0.05 level, of which 61 were within 1Mb of a PD significant SNP. Five genes (*WDR6, CD38, GPNMB, RAB29, TMEM163*; **Table 1**) replicated across Coloc and TWAS results (**Figure 1A**). An additional 5 genes falling outside 1Mb of a PD significant SNP showed strong association for colocalization in both Braineac and GTEx and were significant in the TWAS analysis. Illustrative regional association plots for the highest PPH4 genes and brain regions in Braineac (*RAB29)* in average expression across all brain regions) and GTEx (*CD38* in Putamen) are in **Figure 2A-B.** Full details for all genes that showed strong evidence for colocalization in either Braineac or GTEx, and full details of all genes significant at FDR 0.05 level in the TWAS analysis are found in **Supplementary Tables 2-3.**

**Table 1.**
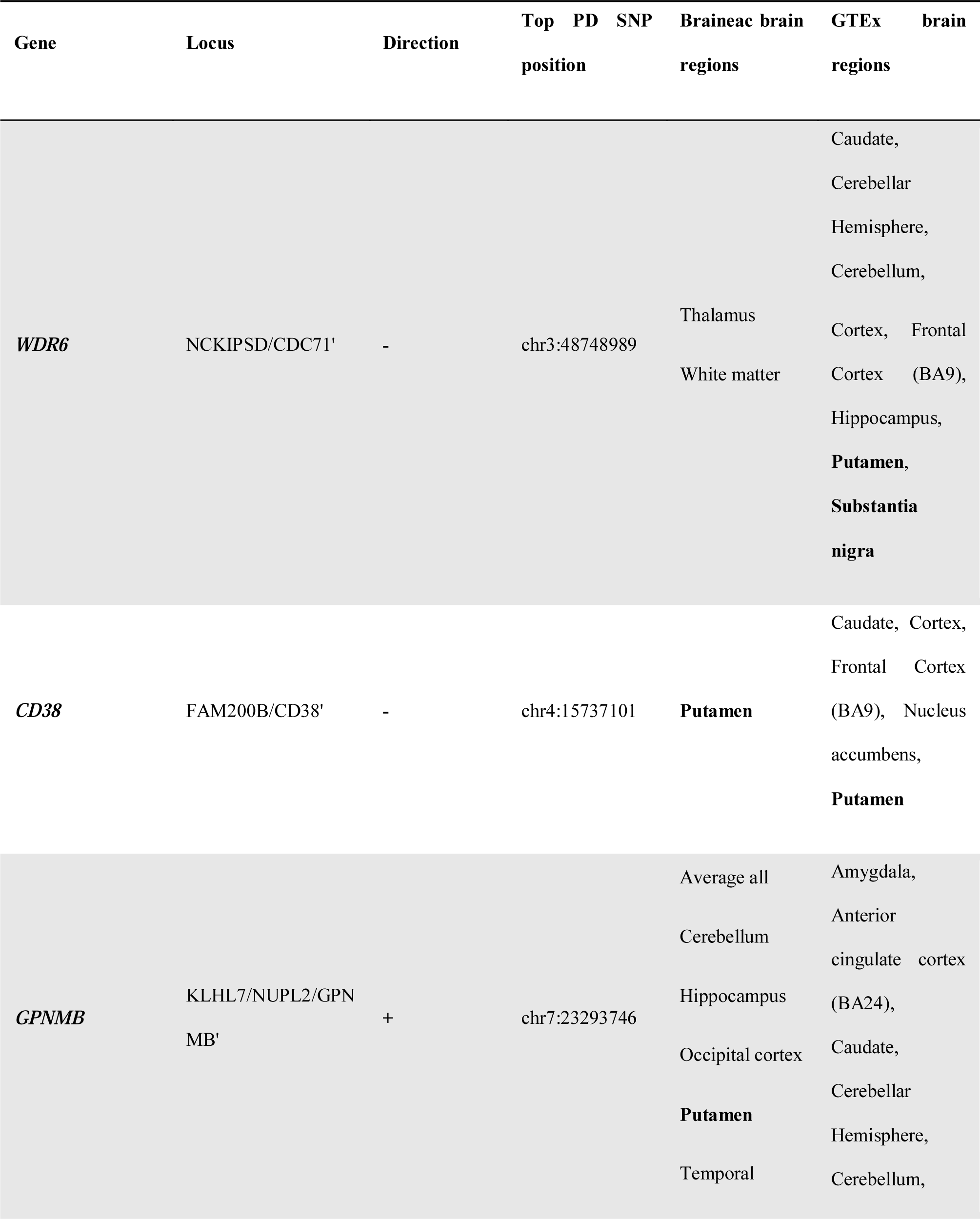
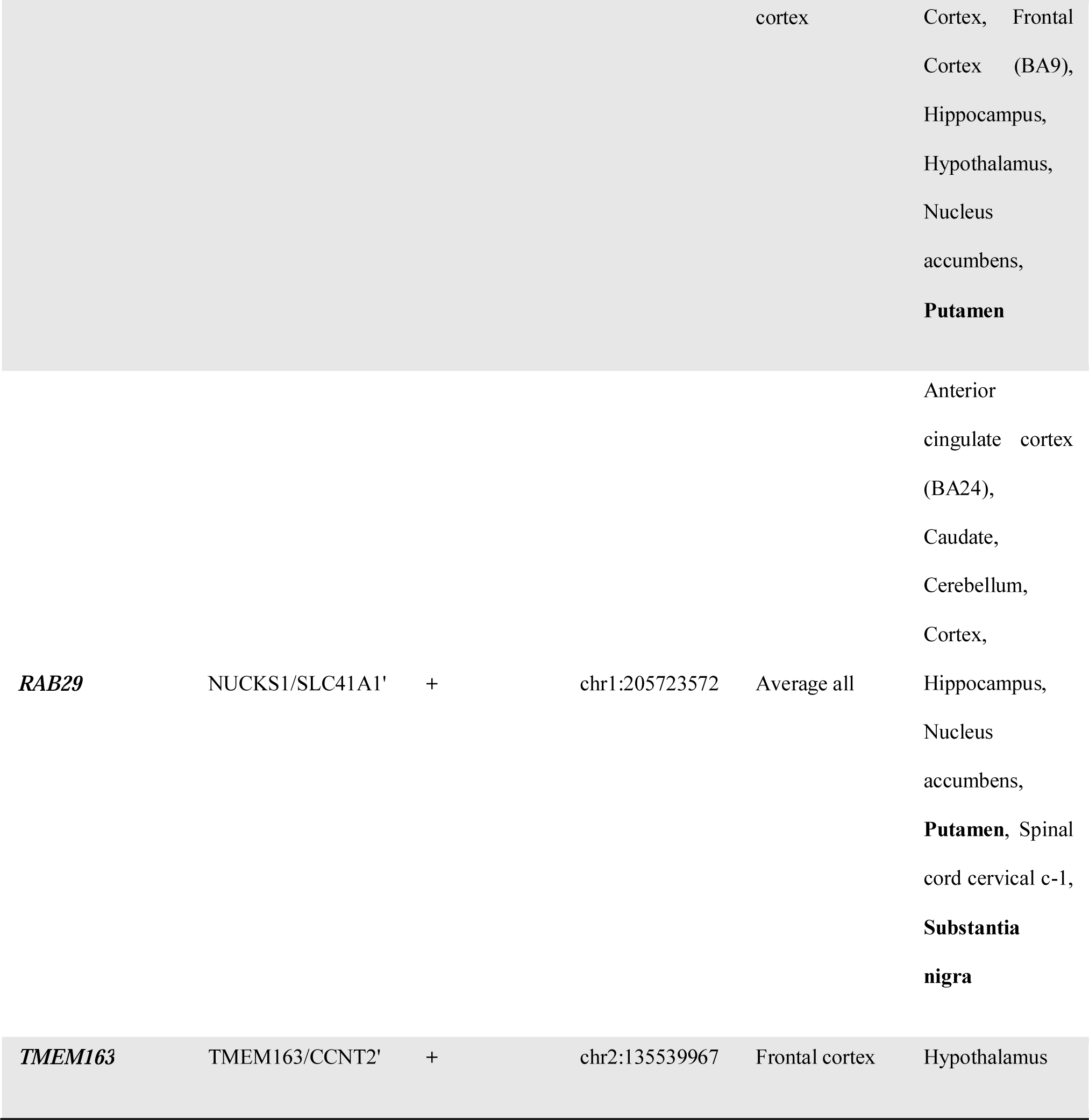
Gene-based results

**Figure 1.**
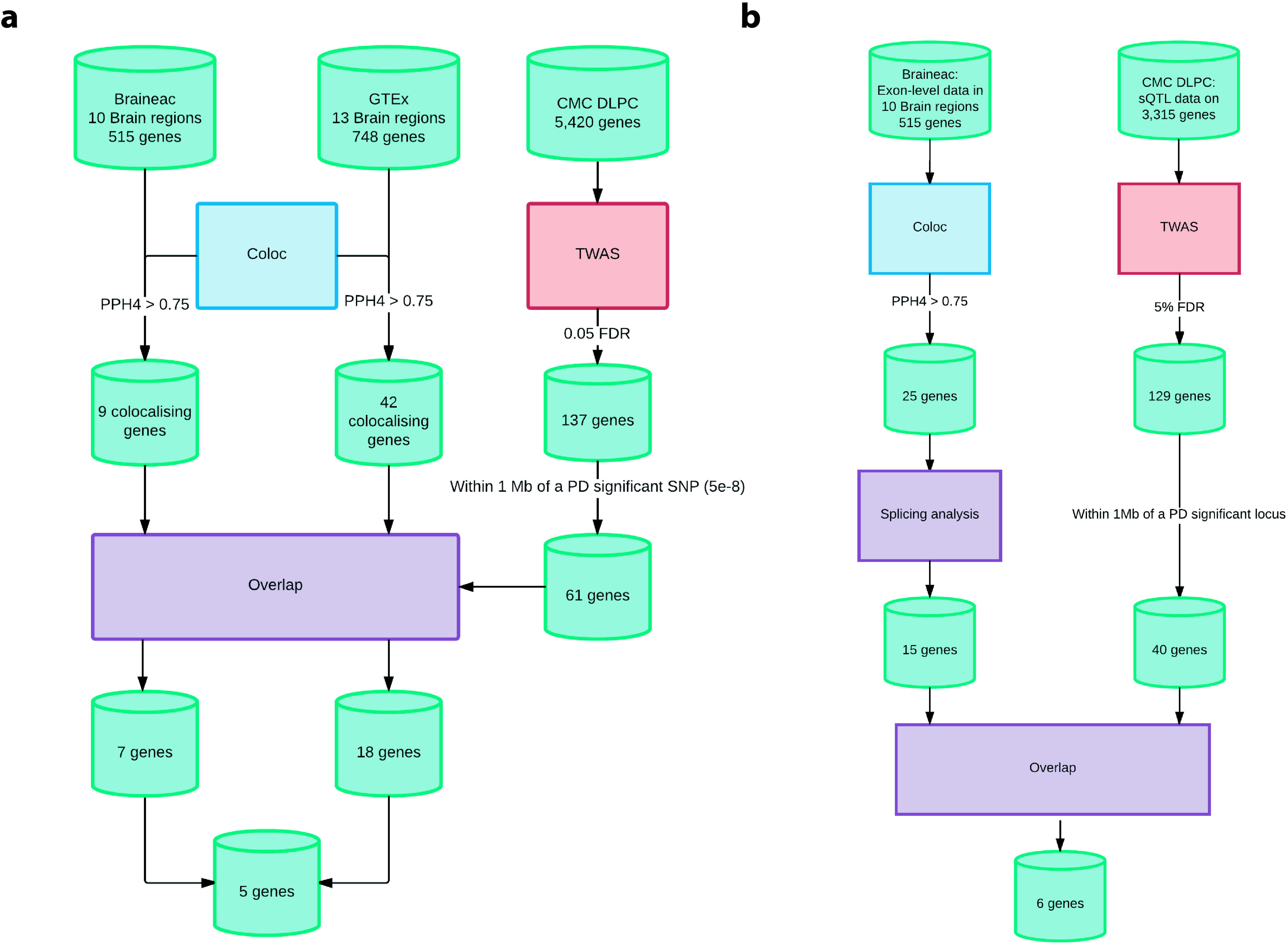
**A:** Flowchart of gene expression analysis. Overall, 5 genes replicated across GTEx and Braineac, and in the TWAS analysis. CMC = CommonMind Consortium; DLPC = dorsolateral prefrontal cortex; TWAS = transcriptome-wide association study. **B:** Flowchart of splicing analysis. Overall, 6 genes replicated across Coloc splicing analysis in Braineac and TWAS sQTL analysis. For Coloc, the splicing analysis consisted of identifying genes with evidence for colocalization at the level of at least one exon (exon PPH4 > 0.75) and evidence against colocalization for the expression of the whole gene (gene PPH3 > PPH4). CMC = CommonMind Consortium; DLPC = dorsolateral prefrontal cortex; TWAS = transcriptome-wide association study.

**Figure 2.**
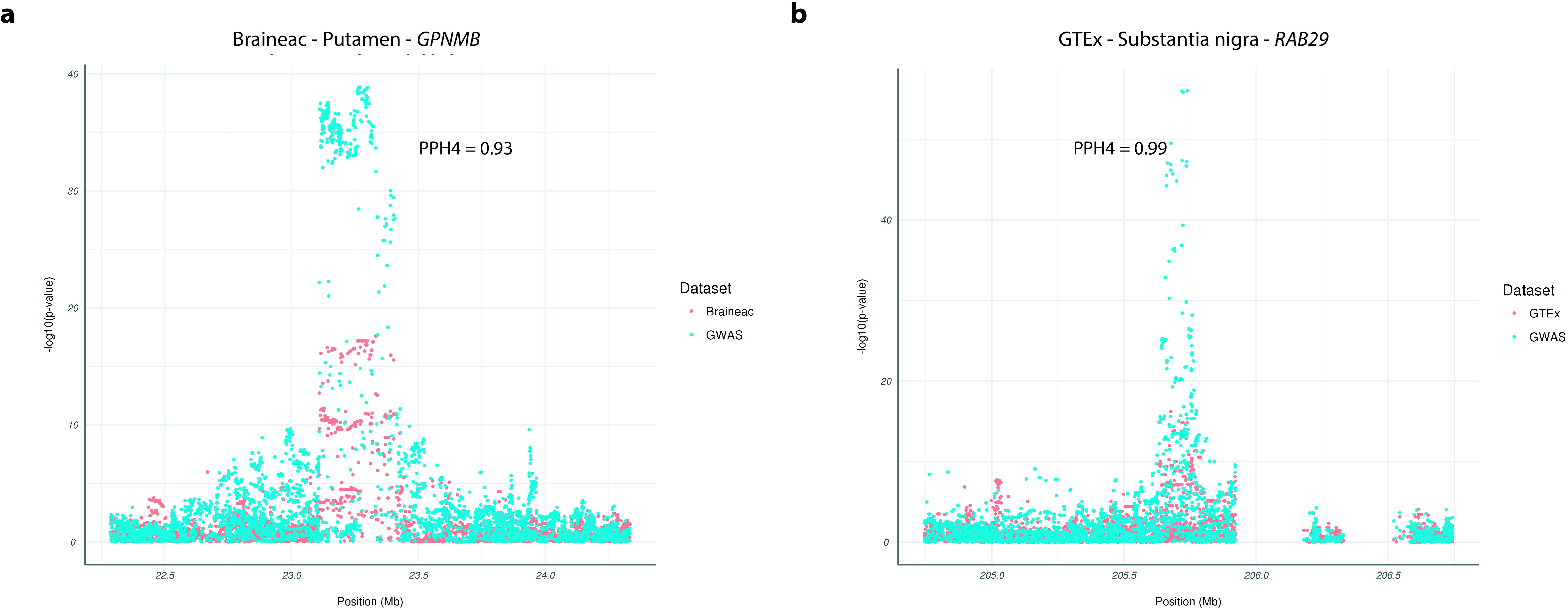
Regional association plots of GPNMB with Braineac data **(A)** and RAB29 with GTEx data **(B)**. The -log10 p-values are presented on the association of each SNP with gene expression (orange) and risk of PD (green), illustrating the likely colocalization of the signals and the likely presence of a shared causal variant.

### Splicing results

Overall, 25 genes had strong evidence for colocalization for at least one exon in at least one brain region in Braineac. For 15 genes there was evidence suggesting that the association is driven by an exon-level splicing event (exon PPH4 > 0.75) rather than a gene-level expression effect (gene PPH3 > PPH4). In the TWAS analysis, 129 genes had evidence for splicing in at least one isoform at FDR 0.05 level. Of these, 40 were within 1Mb of a PD significant SNP. 6 genes with a putative splicing effect in the Coloc analysis showed a significant splicing effect in the TWAS analysis (*ZRANB3, PCGF3, NEK1, NUPL2, GALC, CTSB*) (**Figure 1B**). We then assessed the eQTL p-values of the top SNP suggested by Coloc for the associated exon and the gene as a whole, showing that for the vast majority of these associations, the gene-level p-value is not significant, while the exon-level is. These genes are summarized in **Table 2**. An additional 2 genes falling outside 1Mb of a PD significant SNP showed evidence of splicing in all datasets and analyses. Full details for all genes that showed strong evidence for colocalization in either Braineac or GTEx, and full details of all genes significant at FDR 0.05 level in the TWAS analysis are found in **Supplementary Tables 5-6.**

**Table 2.**
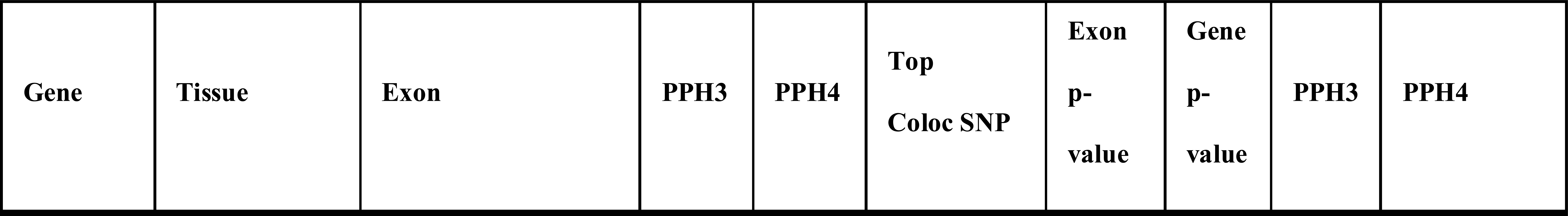
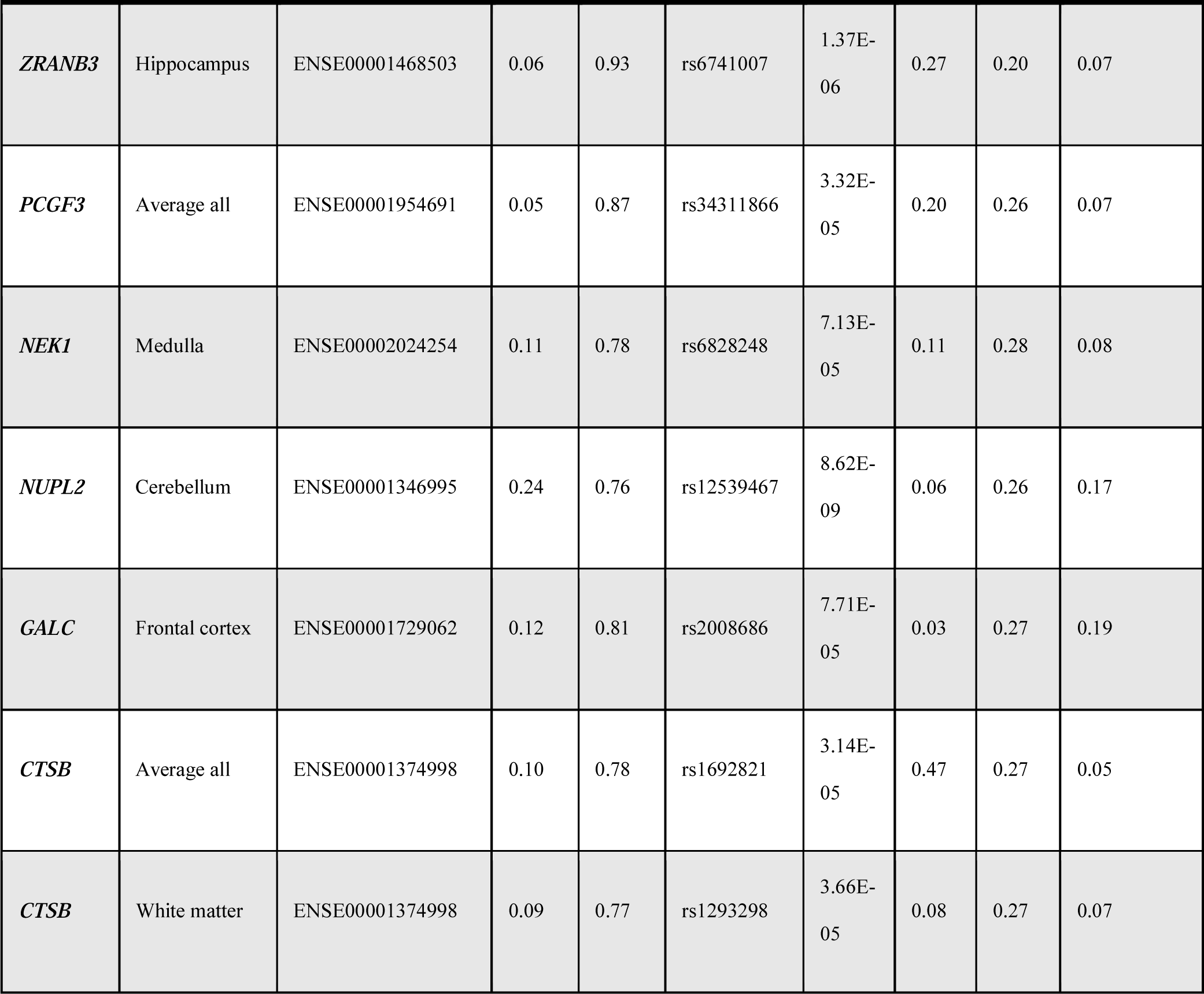
Splicing results

### MWAS results

We then integrated PD summary statistics with summary level methylation data for 37,460 CpGs. 134 CpGs survived FDR correction and conditional analysis in substantia nigra (mapping to 107 unique genes) and. 117 CpGs survived FDR correction and conditional analysis in substantia nigra (mapping to 93 unique genes) (**Supplementary Table 6**). Of the MWAS significant genes, 3 overlap with the Coloc expression or splicing hits (**Table 3**).

**Table 3.**
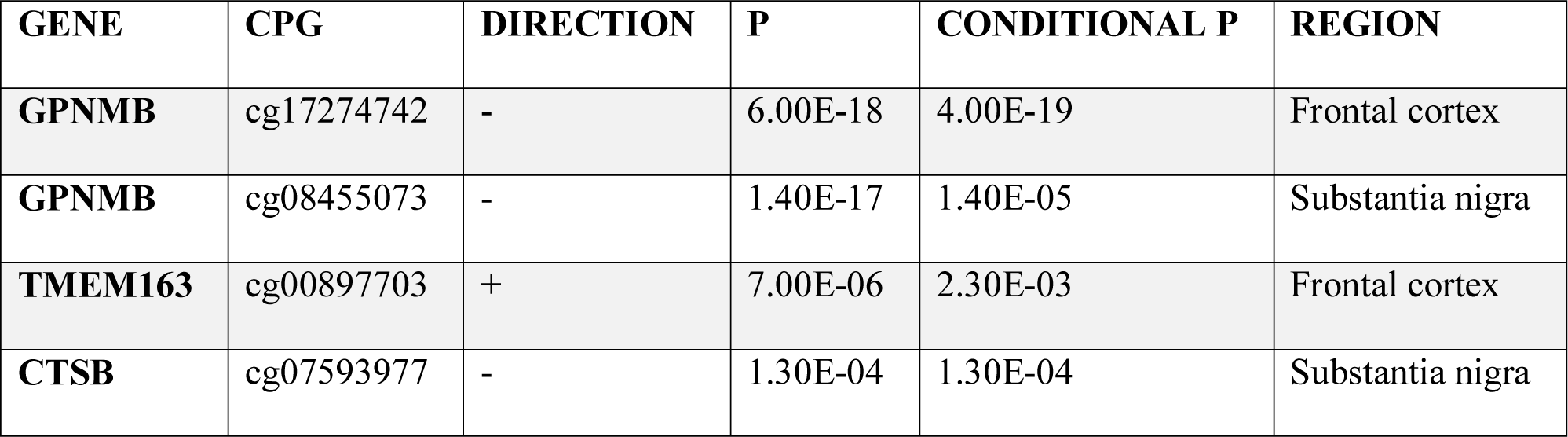
MWAS hits overlapping with Coloc hits

### Cell-type specificity and WGCNA

The results for the cell-type specificity analysis are shown in **Figure 3**. Although no single cell-type dominated, Coloc prioritized gene expression was overall more prevalent in glial cell-types compared to neurons. This finding was consistent across analyses performed using mouse immunopanning data generated from cortex, human-immunopanning data also generated using cortical tissue and using inferred cell-specific gene expression generated using co-expression networks across all brain regions including substantia nigra. WGCNA results are summarised in **Table 3**: *NUPL2, TMEM163* and *ZRANB3* were the most relevant genes (MM > 0.76) within 3 modules in different brain regions. *NUPL2* was a key gene within the darkturquoise, blue and skyblue modules in *nucleus accumbens, caudate* and *putamen* respectively. Interestingly, these modules’ most relevant functions indicated catabolic processes related with protein ubiquitination (protein ubiquitination [GO:0016567, p=7.47e-10]; ubiquitin-dependent protein catabolic process [GO:0006511, p = 2.57e-17]). *TMEM163* and *ZRANB3* were both important in the turquoise module in the *frontal cortex* and *caudate* respectively. This module indicated chemical transmission at the synapse as a major associated function (regulation of signalling [GO:0023051, p=1.33e-65]; cell communication [GO:0007154, p=7.55e-35]).

**Figure 3.**
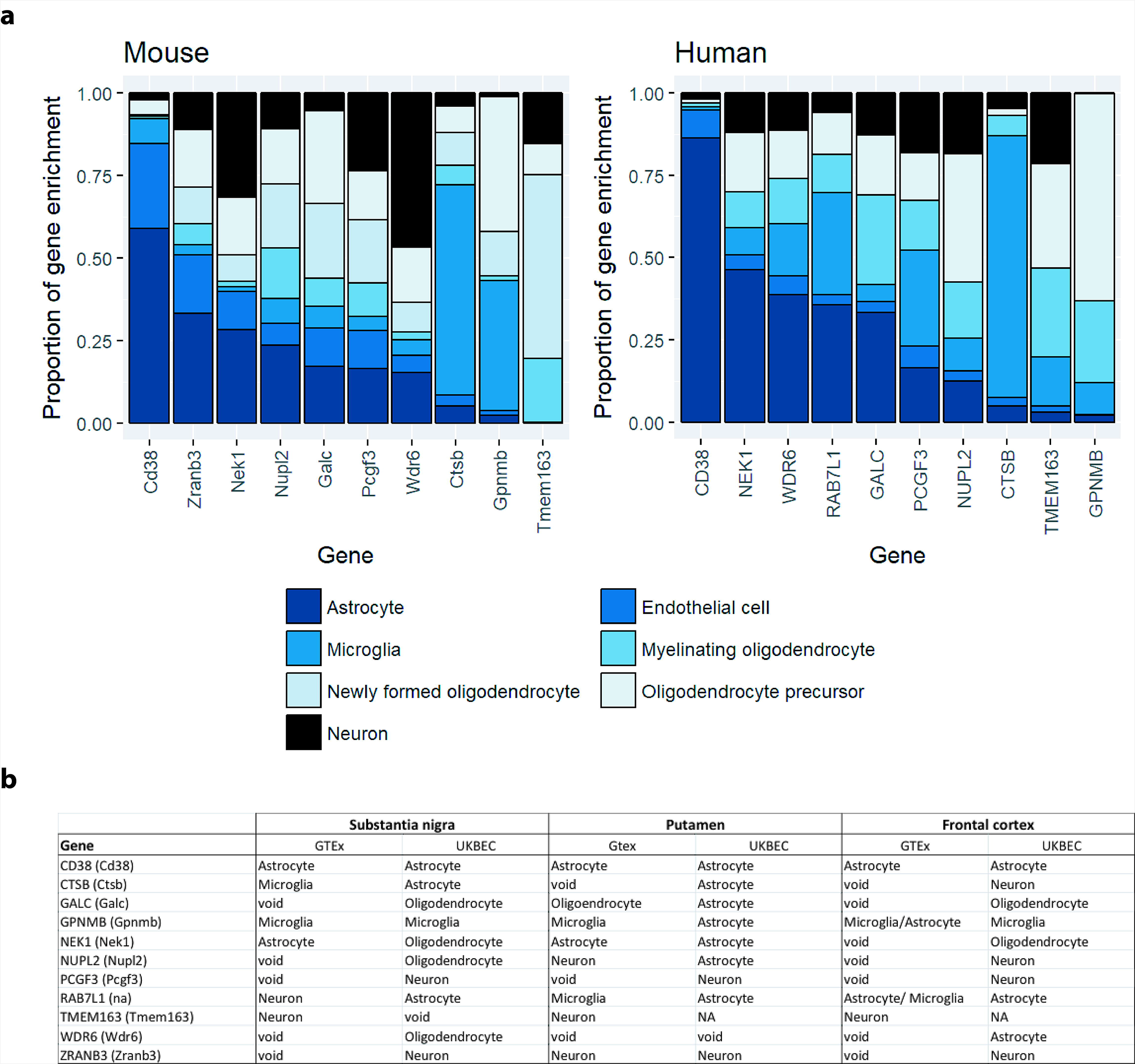
Cell-type specific expression of Coloc prioritised genes in human and mouse, and using GTEx and Braineac data. These results illustrate the overrepresentation of glial cell types compared to neuronal cell types among the candidate genes.

### Literature-derived PPI network

Interestingly the protein products (WPPINA analysis) of the 11 genes prioritized by the Coloc analysis revealed connection (principally through a second degree level of connection) with a number of proteins that are also relevant for Mendelian forms of PD and parkinsonism (**Figure 4A**). The number of connections (n=9) was higher than expected based on random simulation of protein connectivity (**Figure 4B**) obtained by analysing the connections of 1000 control networks characterized by the same number of seeds as the experimental network. Control networks were built by using random combinations of seed-genes sampled out of the pool of 118 genes characterized by evidence against colocalization within the Coloc analysis (PPH3>0.75). Most of these random networks resulted in no connections with PD/parkinsonism related proteins, with the maximum observed number of connections of 4. This result suggests a disease specific and consistent interaction between protein products of the Coloc genes and Mendelian PD/parkinsonism proteins.

**Figure 4.**
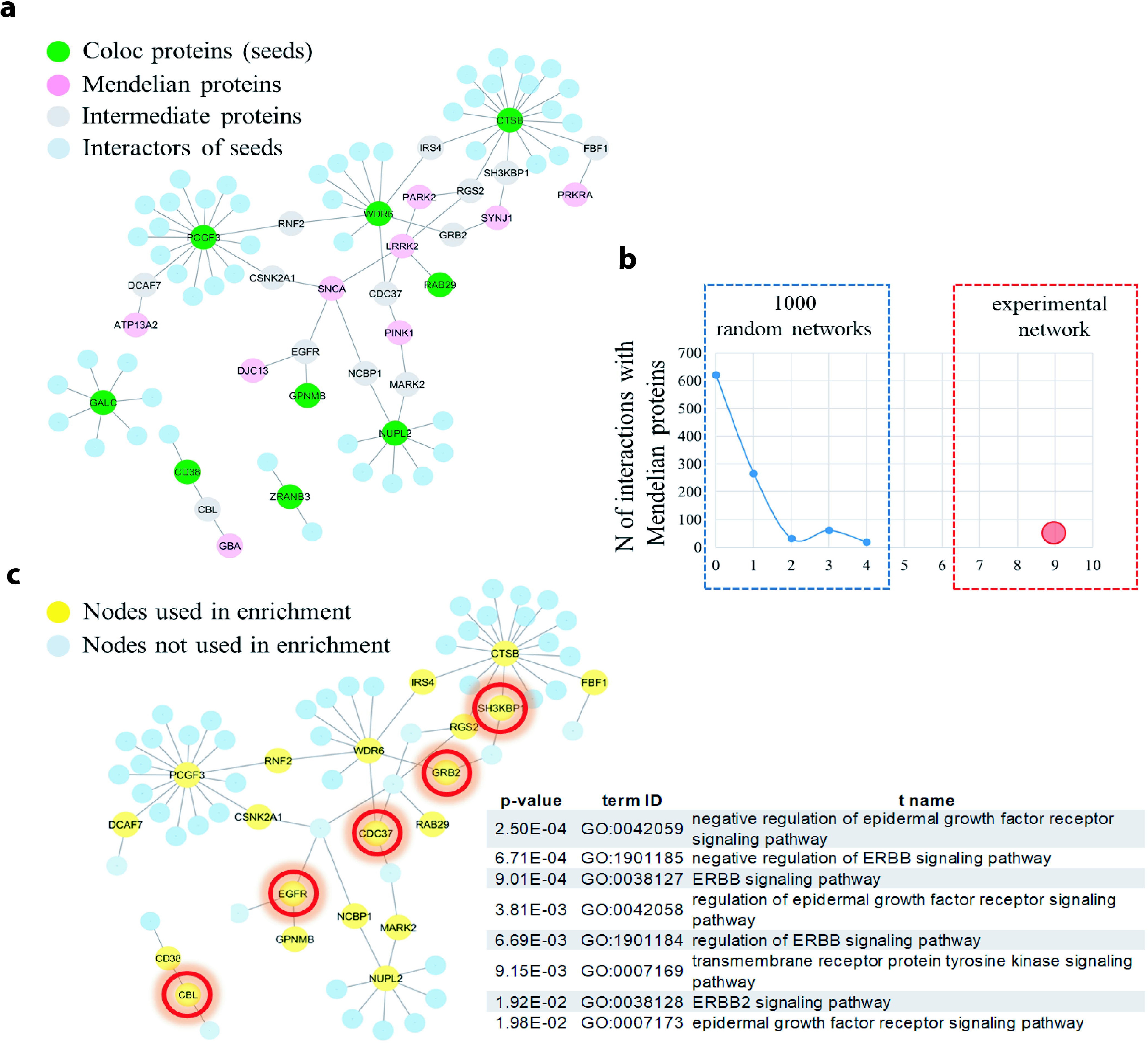
Literature-derived PPIs network. (A) WPPINA network visualisation of the PPIs specific for the proteins (seeds) coded by the Coloc genes (green nodes). Minor protein interaction partners are shown in blue, whilst Mendelian Parkinson’s/parkinsonism proteins interacting with parts of the seeds’ interactome are reported in pink. Major interaction partners, i.e. that they bridge interaction between at least a Coloc protein and a Mendelian protein are labelled in grey. (B) The negative control protein network has been randomly sampled to generate 1000 random networks with similar features to the actual Coloc network. These therefore included same/similar number of seeds (9 seeds) to the Coloc protein network and were matched to the Mendelian protein network to quantify the number of Mendelian proteins able to interact with the random seeds’ interactome. Random simulation and experimental results are shown in blue and red, respectively. (C) Nodes highlighted in yellow (Coloc proteins connected to Mendelian PD proteins and internodes) were used to run functional enrichment. The most specific terms of enrichment are reported in the table with their adjusted p-value, GO term identifier and name. The proteins that contributed to the enrichment of the terms reported in tables are circled in red.

Since proteins that interact together are likely to share similar functions, we investigated if there were shared pathways/biological processes associated with this core of connected Coloc and Mendelian proteins (input list for enrichment in **Supplementary Table 8**). Results are summarized in (**Figure 4C**) and reported in **Supplementary Table 9**, and suggest that there is an enrichment of proteins involved in or regulating the ERBB receptor tyrosine protein kinase signalling pathways.

## Discussion

With the increasing number of GWA studies, our ability to map disease associated variants exceeds our ability to interpret their biological function. Here, we have performed a comprehensive analysis by colocalization of eQTL and GWAS signals in PD, using the latest PD-GWAS. We have interrogated these data with publicly available brain eQTL datasets (Braineac and GTex). This multi-layered approach has identified 11 genes, which we postulate underlie PD risk. Of these, 5 affect gene expression regulation and 6 influence alternative splicing. We found evidence of methylation regulation for 3 of these candidate genes.

Of the candidate genes presented here, *CD38* is involved in insulin regulation, emphasizing a possible role of glucose metabolism in PD (Okamoto *et al.*, 1997). This is supported by recent work indicating a relationship between BMI and PD, and a randomized controlled trial of exenatide, a glucagon-like peptide-1 receptor agonist, as a disease modifying agent for PD (Athauda *et al.*, 2017; Noyce *et al.*, 2017). Furthermore, a role for *CD38* in regulating neuroinflammation, especially in glial cells, has been proposed, which is consistent with the enrichment of *CD38* in astrocytes in the cell-type specificity analysis (Wei *et al.*, 2009). The link between *GALC* and Krabbe adds weight to the recent work on the role of lysosomal pathways in PD (Wenger, 2011; Dehay *et al.*, 2013; Robak *et al.*, 2017). Furthermore, the data from this study reinforces that status of the *RAB29* gene as the priority candidate for the chromosome 1q32 locus association. Recent studies providing further functional evidence linking *RAB29* (aka *RAB7L1*) to *LRRK2*, and implicating *RAB29* as a substrate for *LRRK2* kinase activity, also support this designation (Fujimoto *et al.*, 2018; Liu *et al.*, 2018; Purlyte *et al.*, 2018). Interestingly, in the *GPNMB/NUPL2* locus, the PD GWAS results suggest only one independent signal, while the results presented in this paper nominate both *GPNMB* (gene level) and *NUPL2* (splicing) with strong PPI evidence connecting both to Mendelian or sporadic risk genes. This could be explained by the true causal gene being one of the two, or a single mechanism mediated through the effects on both genes, or potentially yet undetected independent PD GWAS signals at the locus affecting independent risk genes.

The WGCNA and WPPINA approaches allow prioritization of genes as a global functional unit. The WGCNA analysis suggested that 3 of the Coloc prioritized genes (*NUPL2, TMEM163* and *ZRANB3*) may be relevant for supporting functions related with the ubiquitin proteasome system, neuronal development and the chemical transmission at the synapse. The WPPINA analysis indicated that the proteins encoded by the Coloc prioritized genes interact with Mendelian PD/parkinsonism proteins suggesting the existence of a common functional unit of genes/proteins – related to the ERBB signalling pathways – that increases the risk for developing sporadic as well as familial PD.

It is noteworthy, that this study has considered only *cis*-eQTLs, due to the current challenges in robustly quantifying *trans*-eQTLs. Furthermore, this study only considered eQTLs in the brain and no other tissues. The Coloc analysis applied here assumes that the true causal variant underlying the disease has been captured in both the GWAS and eQTL datasets. The PD-GWAS data used here has been imputed to the latest HRC panel (v1.1), and the genotypes in the GTEx data are generated with whole-genome sequencing, maximising the chances of this assumption being met. However, the Braineac data used here has been imputed to 1000 Genomes phase 1, potentially reducing our power to replicate candidate genes in this dataset. Another limitation in the colocalization tools used here is that they assume one independent signal for each gene at each locus for both the GWAS and QTL results. Finally, the methods used here cannot exclude pleiotropy, whereby a disease-causing SNP affects the regulation of an unrelated gene via a separate pathway.

While the overlap between GTEx and Braineac derived results are encouraging, there are some inconsistencies. This may be due to potential methodological differences in tissue collection, RNA extraction, platforms, and analyses pipelines. in addition, these differences might reflect divergent cell-type specificity of the expression and splicing effects.

A key strength of this study is that this is a large and comprehensive exploration of PD GWAS and eQTL dataset from human brain. We replicated our Coloc results across two platforms: Braineac and GTEx, generated through microarray, and RNA-seq, respectively, and performed additional validation using TWAS in the CMC DLPC dataset. This has resulted in prioritizing 11 candidate causal genes for PD based on GWAS hits to be further investigated biologically in different animal or cell models for PD. Furthermore, we are highlighting biological reasons for their likely functional contribution to PD pathogenesis. We acknowledge further functional work will be required to mechanistically link these genes to PD, but the genetic and analytical approaches applied here suggest that these are the putative gene and genomic events underlying these risk loci. Techniques such as chromosome conformation capture and generating cell models with altered gene expression and splicing patterns will be key to characterise the potential role of these genes in PD pathogenesis.

## Acknowledgements

The Braineac project was supported by the MRC through the MRC Sudden Death Brain Bank. P.A.L. was supported by the MRC (grants MR/N026004/1 and MR/L010933/1) and Michael J. Fox Foundation for Parkinson’s Research. D.T. was supported by the King Faisal Specialist Hospital and Research Centre, Saudi Arabia and the Michael J. Fox Foundation for Parkinson’s Research, J.H acknowledges support from the Dolby Foundation. R.F. is supported by Alzheimer’s Society (grant number 284). University College London Hospitals and University College London receive support from the Department of Health’s National Institute for Health Research (NIHR) Biomedical Research Centres (BRC). NWW is an NIHR senior investigator and receives support from the JPND-MRC Comprehensive Unbiased Risk factor Assessment for Genetics and Environment in Parkinson’s disease (COURAGE). R.H.R is supported through the award of a Leonard Wolfson Doctoral Training Fellowship in Neurodegeneration.

